# Doctoral Students as Carbon Accountants: Calculating Carbon Costs of a PhD in Neuroscience

**DOI:** 10.1101/2025.01.20.633775

**Authors:** William V. Smith, Aimee Bebbington, Ranjini Sircar, Stefan R. Pulver

## Abstract

Research is an energy and resource-demanding activity. However, despite increasing awareness and emerging sustainability initiatives, a paucity of data and methodological inconsistency continue to hamper effective and accountable emissions mitigation. With > 250,000 doctoral students graduating annually across all academic disciplines, empowering PhD students to engage in carbon accounting could provide a sizable and robust source of carbon data alongside a powerful generational force for decarbonisation. Here, we demonstrate how doctoral students and other researchers can consistently measure the carbon footprint of their work, using one PhD student’s research in a neuroscience *Drosophila* lab as our case study. We present a comprehensive life-cycle assessment of the equivalent carbon dioxide emissions (CO_2_e) generated by the student’s research activities, including measurement of scope 1 emissions associated with *Drosophila* husbandry; calculation of time- and region-specific scope 2 emissions produced by widely used techniques including calcium imaging, electrophysiology, and optogenetics; and estimation of scope 3 emissions associated with procurement and research-related travel. We found that research-related travel and procurement of laboratory supplies were responsible for the majority of annual emissions, up to 1942 kg CO_2_e and 543 kg CO_2_e respectively after accounting for aircraft radiative forcing. Using NESO’s open-source Carbon Intensity API to account for temporal and geographical variation in the carbon intensity of UK National Grid energy, we found that persistent laboratory energy consumption released 10.99 kg CO_2_e, with an additional 3.56 kg CO_2_e scope 2 and 3.6 kg CO_2_e scope 1 emissions underpinning direct research activities. Finally, we discuss the challenges of accurately carbon foot printing research across disciplines in the UK and beyond, highlighting the value of regionally precise open-source energy mix data and the need for data openness within research supply chains. Overall, we present a common framework for including carbon footprint analyses as ‘Carbon Appendices’ to PhD theses to generate carbon footprint data across disciplines. Beyond the benefits of such data for informed emissions mitigation, we envision doctoral students carrying insights from carbon appendices forward into academia and industry to catalyse a community-driven decarbonisation of the research sector.

## Introduction

Greenhouse gas emissions resulting from human activities are a major driver of global warming and climate change. Efforts to systematically reduce and mitigate the effects of excess atmospheric greenhouse gases involve coordinated actions across all sectors of life. Scientific researchers are increasingly considering the environmental impacts of their work. By focusing on “cradle-to-grave” life cycle assessments, institutions (McGain and Naylor, 2014; Valls-Val and Bovea, 2021; Martin *et al*., 2022) and individual researchers (e.g., Thiel *et al*., 2017; Gordon *et al*., 2021; Elli *et al*., 2024) have begun to account for their direct (scope 1) and indirect (scope 2, via energy production; and scope 3, incorporating all other life cycle emissions not operated or controlled by the user) activity-related carbon emissions, with the goal to mitigate lab activity, procurement, and disposal-related carbon costs. At the individual level, researchers have contributed to estimating the carbon associated with specific resources (e.g., Montoro *et al*., 2023) and domain-specific practices (e.g., Grealey *et al*., 2022). At the institutional level, organisations across the world have devised practices and reported on the carbon footprint of sectors including healthcare (Purohit, Smith and Hibble, 2021; Robinson *et al*., 2023; Maida *et al*., 2024) agriculture (Mazzetto, Falconer and Ledgard, 2022; Ozlu *et al*., 2022), and industrial manufacturing (Panagiotopoulou, Stavropoulos and Chryssolouris, 2022). Individual-led action in computation (Lannelongue, Grealey and Inouye, 2021), and clinical sciences (Farley and Nicolet, 2023) has demonstrated the power of researchers in generating carbon footprint data from a variety of specialised domains. Such actions have supported collective action by researchers to design accredited sustainable waste disposal initiatives (e.g., LEAF and MyGreenLab) (Schell and Bruns, 2024). However, decentralised efforts are in their infancy in many disciplines especially beyond clinical medicine and agriculture, with partial reporting on equipment and techniques alongside methodological inconsistency hampering effective carbon footprint estimation and weakening cohesive and coherent mitigation strategies (Rae *et al*., 2022; Robinson *et al*., 2023).

Scientific research is one domain of the carbon economy that has been underexplored and may come to increasingly dominate the landscape of energy usage in the coming decades. In particular, as science progresses, the complexity of the problems we address rises. The advancement in high-energy experimental techniques, the exponential rise of, and dependency on, artificial intelligence in analytical pipelines, and the increasing accessibility of science throughout the world means the carbon burden generated by science will continue to rise as we approach our climate targets (Budennyy *et al*., 2022; Tamburrini, 2022). Understanding the current carbon footprint of science is hampered by partial reporting with major gaps in our awareness of costs across the life cycle of research (i.e., procurement, techniques, disposal). While scientific institutions are all directed to the same goal of knowledge creation, the different methodologies for scientific research - specialisation of equipment, techniques and protocols, recycling methods, mitigation strategies - can vary significantly. Consistent internal assessments of carbon footprints by researchers worldwide could provide an efficient, sustainable path to building an accessible data set and attentive accountants of discipline-specific carbon footprints, to focus and improve institutional, governmental, and industry decarbonisation efforts.

There are many methods for estimating carbon footprints, with varying complexity, which can make individuals and institutions reluctant to engage in carbon accounting (Fenner *et al*., 2018; Robinson *et al*., 2018). One solution is to create a cross-discipline community utilising the same methodologies to consistently report research-specific carbon footprints. To enable this, we propose a framework to quantify emissions emerging from individual PhD theses. We hope that these quantifications can contribute to data on, and serve as interim proxies for, the emissions of multiple disciplines and institutions. There were 115,705 PhD students studying in the UK in 2022/4 (HESA, 2024), with an estimated 277,700 doctorates awarded annually worldwide (OECD, 2019) Many of these students will become part of the next generation of academics and public servants; thus, engaging doctoral students with carbon accounting could help strengthen commitment to sustainability within academia and industry while generating a large volume of reliable data for effective and accountable carbon mitigation strategies. Furthermore, each student generates a thesis which can serve as a common vehicle for disseminating information about the carbon footprints of specific research activities.

Here, we demonstrate how PhD students can consistently measure the carbon footprint of their work, using one doctoral student’s research in a neuroscience *Drosophila* lab as our case study. We present the carbon costs associated with each stage of the research life cycle, from procurement through to experiments, analysis, data storage and material disposal. As part of this life-cycle assessment, we calculate estimates of the carbon cost associated with many activities relevant to other neuroscience and *Drosophila* labs, including scope 1 emissions from anaesthetisation of flies; scope 2 emissions from food preparation, commonly used experimental equipment and analytical pipelines, and waste disposal; and scope 3 emissions associated with fly requisitions from stock centers. Additionally, we calculate the total carbon cost associated with one year of PhD work in neurophysiology (including research and travel), using robust, easily replicable methodologies. Finally, we explore reported variations in the equivalent carbon dioxide emissions (CO_2_e) associated with UK National Grid energy to better understand the role of geography and time in the production of scope 2 emissions from research. Overall, we demonstrate how PhD students and other researchers can begin to account for the carbon costs of their work and discuss how uptake of this approach could improve research sustainability by informing targeted emissions mitigation strategies.

## Results

### The Carbon Cost of *Drosophila* Neuroscience Research

Reliable carbon accounting in academic research is vital for tailoring effective mitigation strategies and generating a continuous stream of robust data to hold implementation policies accountable. *Drosophila* neuroscience research generates scope 1 (direct), 2 (indirect via electricity production), and 3 (incorporating all other life cycle emissions not operated or controlled by the user) carbon emissions. Over the course of one year, we conducted life cycle analyses that accounted for the carbon generated by all stages of one PhD student’s (the first listed author’s) research activities, assuming all energy came from the mains network grid in Southern Scotland. Here, we detail the measured and estimated carbon dioxide equivalent emissions (CO_2_e) for each stage of the research life cycle in a chronological manner commencing with food generation and finishing with material disposal and data storage (Figure 1A). We describe how we quantify carbon emissions from all stages of the research cycle. A summary of this quantification is provided in Figures 1B and 1C.

**Figure 1:**
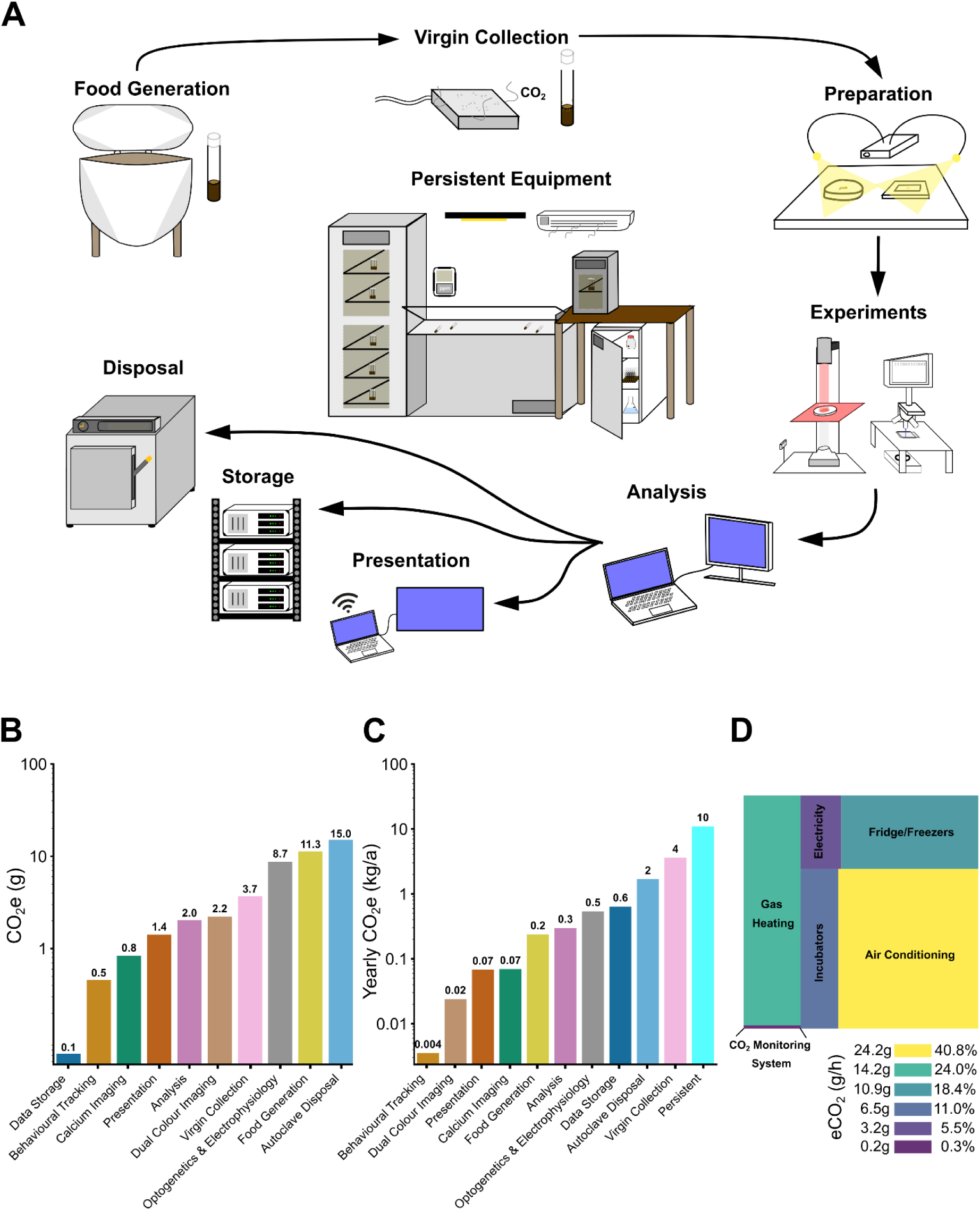
The carbon costs associated with one year of *Drosophila* neuroscience research. (**A**) The life cycle of *Drosophila* experimental research. (**B**) Equivalent carbon dioxide emissions, CO_2_e (g), per experimental preparation (behavioural optogenetics, calcium imaging, electrophysiology), per preparation (analysis), per vial (virgin collection), and per 700 medium-size vials (food generation, disposal), to 1 d.p. (**C**) Yearly equivalent carbon dioxide emissions CO_2_e (kg/a) generated from the work of one researcher in South Scotland for June 2023 - May 2024 including all aspects of the research lifecycle based on total number of experiments completed, vials used, and data generated per month. (D) The average hourly equivalent carbon dioxide emission rate, CO_2_e(g/h), calculated from thepower drawn by different persistent laboratory equipment and the average monthly kWh-to-CO_2_e conversion factor for Southern Scotland.

#### Food generation

All animal research depends on food generation. This often relies on in-house equipment, which generates scope 2 carbon emissions via electricity consumption. For *Drosophila*, food sources consist of a ready-made cornmeal-based resource which is rehydrated, heated and mixed (Lakovaara, 1969) Flies are raised on this medium in vials, with a new generation of flies transferred to fresh vials each month to maintain the genetic stock. In our laboratory, 216 stocks are maintained, and additional vials are used to rear flies for crosses and experiments. Every month, these activities for our research group consumed a combined 700 medium-sized vials with an average of 10g of yeast extract food per vial. The individual PhD student in this study used on average 52 vials per month, amounting to 7.4% of the total laboratory consumption (see Methods). In common with many laboratories, large batches of food are supplied across multiple research groups. Consequently, technicians rely on large-scale, high-power equipment whose energy supply is not easily accessible. Modern research kitchen equipment often incorporates digital monitoring of energy consumption. However, in our case, the use of older equipment precluded direct determination of energy usage, for safety reasons. Thus, we accounted for the maximum electrical power required for food generation based on manufacturer reporting. For heating and mixing of food, we found a maximum power usage of 7.536 kWh (Table 1). We estimated the associated greenhouse gas emissions from electricity generation using region- and time-specific kWh-to-CO_2_e conversion factors. The kWh-to-CO_2_econversion factor varies across time and space due to changes in the balance of power derived from different types of power sources (Raugei, Kamran and Hutchinson, 2020) contributing dynamically to the UK National Grid. To find kWh-to-CO2 conversion factors for Southern Scotland, we used the Carbon Intensity API, a free resource that provides region-specific information on the carbon intensity of UK National Grid energy based on machine-learning forecasting (api.carbonintensity.org.uk). Using estimates of monthly average conversion factors (see Methods), we found that food generation for the whole research group emits on average 152.4 g CO_2_e per month, which, given monthly variations, totalled 3.2 kg CO_2_e for the June 2023 - May 2024 period. Given that our individual PhD student utilised 7.4% of total monthly vials, food generation for this single PhD student’s work emits on average 11.3 g CO2e per month (Figure 1B), which, given monthly variations, totalled 237.8 g CO2e (Figure 1C) for the June 2023 - May 2024 period.

**Table 1:**
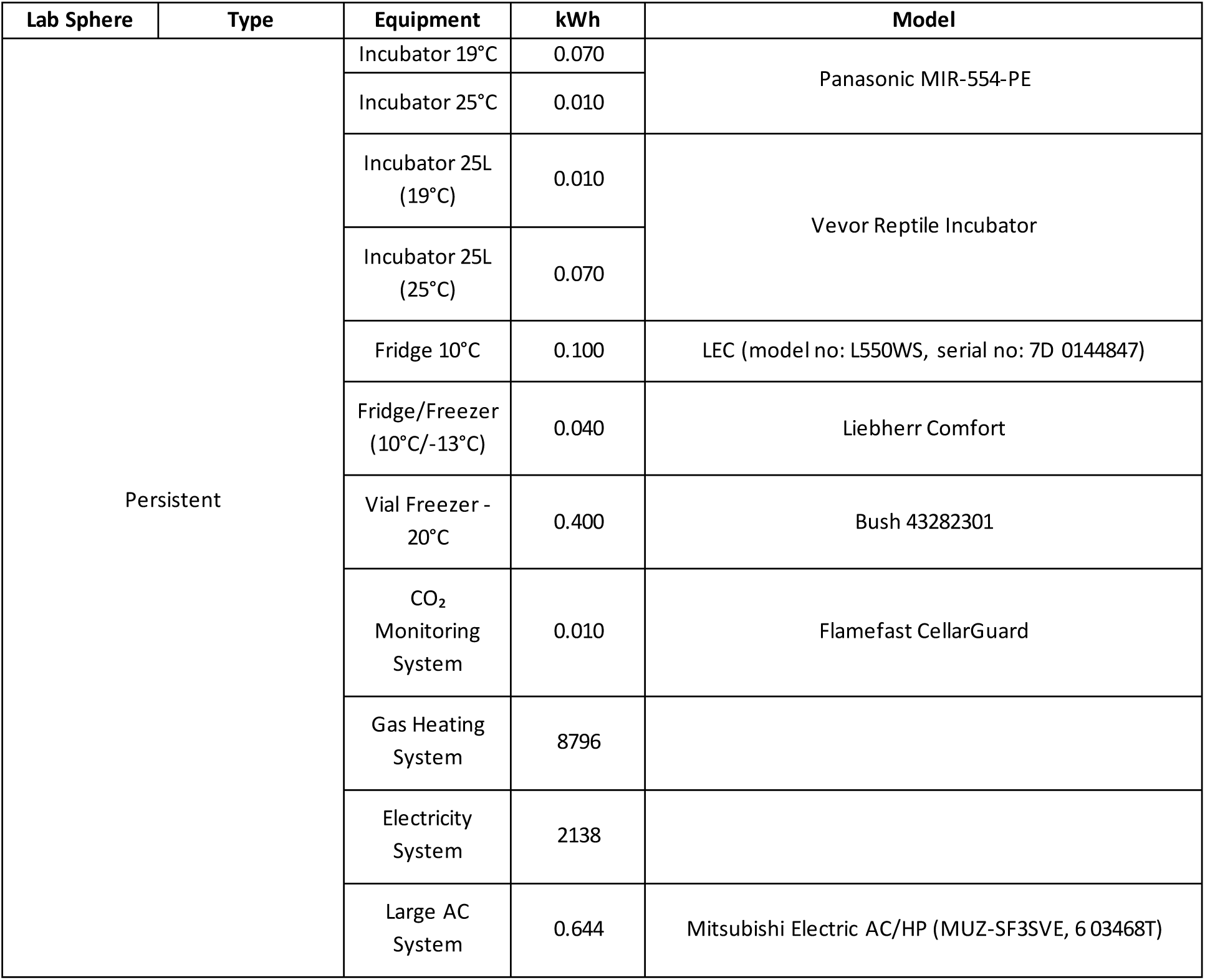
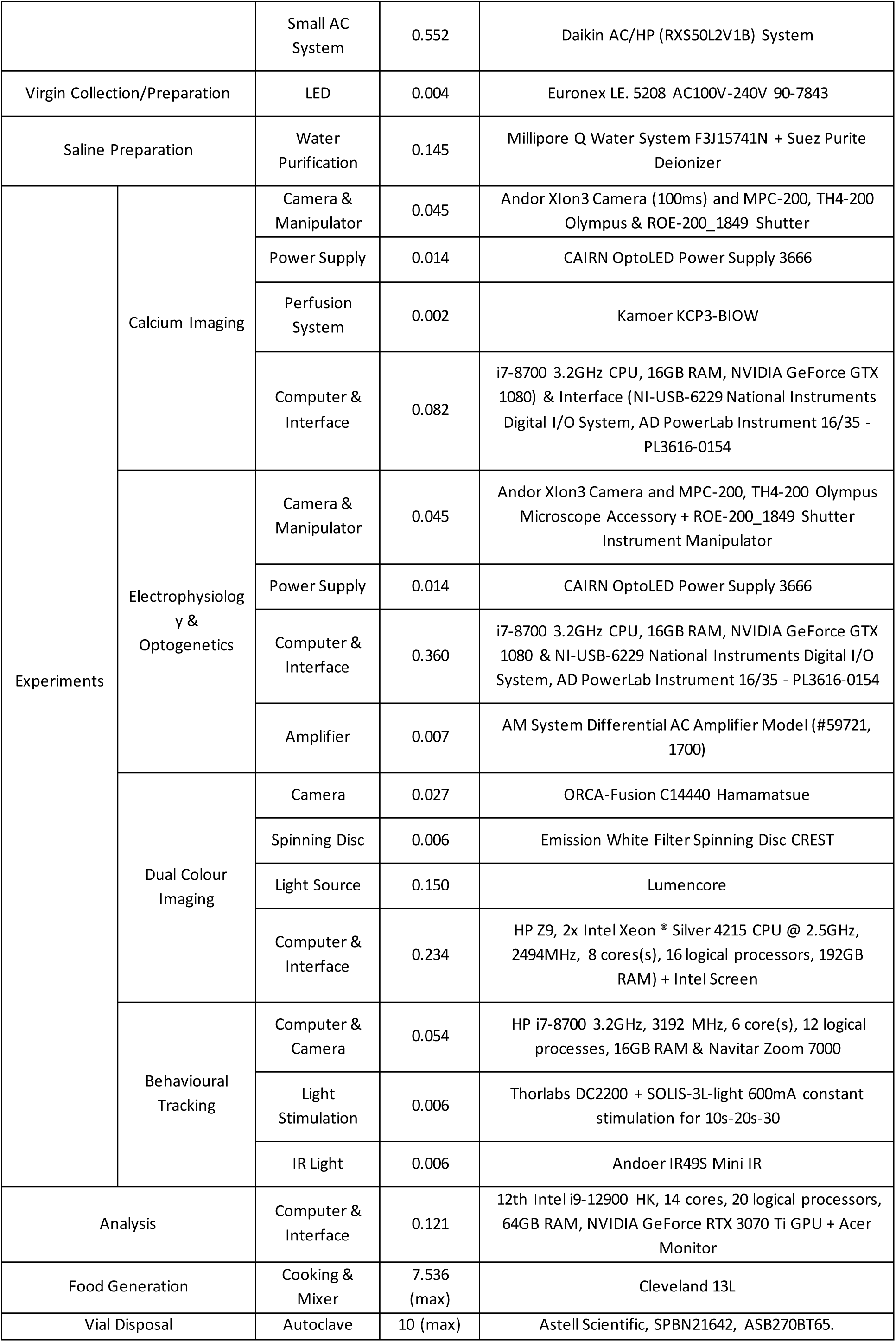
Energy Costs and Model Types for Persistent and Experimental Equipment.

#### *Drosophila* genetic engineering (virgin collection)

*Drosophila* are commonly used model organisms in neuroscience due to ease of genetic manipulations involving selective breeding between transgenic lines. A vital part of this process is anaesthetisation of flies with CO_2_ before sorting newly-eclosed female virgins for genetic crosses, resulting in scope 1 carbon emissions. Anaesthetisation involves the variable use of a CO_2_ gun (up to 2s per vial) and pad (up to 10s per vial). Using a gas syringe and gas displacement method (see Methods), we accounted for the direct release of 3.7g CO_2_ per vial of sorted flies (up to 0.2ml CO_2_). Considering the vast number of laboratories dedicated to *Drosophila* genetics, the level of direct release of CO_2_ from anaesthetisation is likely large but currently unknown. For one PhD student, we calculated a total of 3.6 kg CO_2_e release (from 972 vials) from virgin collection per year (Table 2). However, *Drosophila* labs that rely more heavily on screening genetics (i.e., involving several, consistently used multi-stage genetic crosses like in developmental-based *Drosophila* research) could conservatively directly release <16.7kg CO_2_e (∼4500 vials) a year per researcher (see Methods). Virgin collection often relies upon a microscope with LED illumination to aid in sex determining the adult *Drosophila*.

**Table 2:** CO_2_e Research Costs for One Researcher 2023-4. The yearly CO_2_e (kg) is calculated on the rolling average of the South Scotland kWh/CO_2_e conversion with per task CO_2_e the average of the year.

| Task | Per Task CO<sub>2</sub>e (g/hr) | Yearly Ns | Time Per Task (hr) | Yearly CO<sub>2</sub>e (g) |
| --- | --- | --- | --- | --- |
| Virgin Collection | 1.60 | 972 | ~ | 1555.20 |
| Preparation | 0.18 | 132 | ~ | 23.76 |
| Behavioural Tracking | 0.40 | 8 | 2.5 | 3.53 |
| Calcium Imaging | 0.80 | 94 | 1.0 | 69.34 |
| Dual Colour Imaging | 2.20 | 33 | 0.3 | 24.00 |
| Optogenetics & Electrophysiology | 8.70 | 16 | 2.5 | 539.23 |
| Lab Task | CO<sub>2</sub>e (g/vial) | Yearly Ns | Time per Task (hr) (max vials) | Yearly CO<sub>2</sub>e (g) |
| Food Generation | 0.21 | 8400 | 1 (400) | 3200.60 |
| Vial Disposal | 1.47 |  | 2 (150) | 2240.00 |

#### Preparation and experiments

Next, we wanted to understand how energy use during experimental work contributes to scope-2 carbon emissions. To estimate the carbon intensity of different experiments throughout the year, we used power meters to measure the power drawn by electrical apparatus during experiments, and applied monthly average kWh-to-CO_2_e conversion factors calculated from Carbon Intensity API (api.carbonintensity.org.uk) data for the period June 2023 to May 2024.

Experiments involving optogenetics, calcium imaging, and electrophysiology are staples of modern research especially in neuroscience (Knot *et al*., 2005; Scanziani and Häusser, 2009; Häusser, 2014). All of these experiments utilise high-powered LED lights, high-frame rate (100 frames per second), supercooled (-70 °C) cameras, and persistent signal processing units. In our work, we used these experimental techniques to monitor the motor activity in isolated *Drosophila* neural tissue using genetically encoded calcium indicators and motor root bursting patterns. In the case of electrophysiology and optogenetics, we used pulses of LED-generated light to induce changes in neural network activity, which we measured with continuously-recording electrodes. For dual-colour calcium imaging, we utilised a spinning-disc confocal microscope to acquire images of neurons simultaneously expressing two different genetically encoded fluorescent calcium indicators.

All imaging and optogenetics experiments were performed on isolated neural tissue immersed in physiological saline and dissected under a dissecting microscope fitted with LEDs. Physiological saline (Baines External Saline, (Marley and Baines, 2011)) required the use of deionisation and filtration equipment that yielded 5.27 g CO_2_e for every 4 L of saline produced. Over the 2023-4 research year, saline generation accounted for 26.37 g CO_2_e (20 L). Our microscope’s LEDs yielded, on average, 0.03 g CO_2_e every preparation (up to 10 minutes per prep) amounting to a precursory amount of 4.07 g CO_2_e for all experimental work (151 preparations). Overall, for dissection alone, we calculated an average annual carbon intensity of 30.44 g CO_2_e (Table 2). Preparation time was directly factored into each experimental estimation.

Based on the average kWh-to-CO_2_e conversion factor for 2023-4, we found the average CO_2_e emissions per preparation for each technique (without preparation): behavioural optogenetics (0.5 g); calcium imaging (0.8 g); dual-colour calcium imaging (2.2 g); and optogenetics and electrophysiology (8.7 g) (Figure 1B). Over the course of the year June 2023 - May 2024, we recorded the quantity (151 experiments) and duration (154.5 hours) of all experimental work performed by one PhD student (Table 2). Again, using monthly average kWh-to-CO2e conversion factors, we found that these experiments produced a total of 0.64kg CO_2_e annually (Figure 1C).

#### Data analysis, storage, and presentation

Next, we evaluated the emissions associated with data analysis, storage, and presentation.

Analysis requires computationally demanding and time-intensive processing pipelines. Within a year, the neuroscience PhD student utilised several different CPU-based analytical pipelines to visualise and interpret recorded data. For calcium imaging, image sequences required positional stabilisation over time, extraction and visualisation of fluorescence changes in regions of interest, and relational analysis across many regions of interest. For electrophysiology, nerve root bursting signals require pre-processing rectification and smoothing followed by extracting signal relationships and quantities like bursting duration. For behavioural tracking, videos are analysed using trained neural networks through the software TrackMate (Tinevez *et al*., 2017) and DeepLabCut (Mathis *et al*., 2018) to extract positional and velocity information over time. For the 2023-4 research year, calcium imaging and electrophysiology were the only analytical pipelines utilised. All analysis required extraction and post-processing with programs like DataView (Heitler, 2007) and custom-built Python scripts. Through tracking the hourly power usage on a laptop during five instances of post-hoc analysis (i.e., pre-processing in a signal processing software using DataView, post-processing and visualisation in Python3), we accounted for an average of 2.0g CO_2_e per preparation, yielding 302g CO_2_e across the year.

All of the data collected were stored within a university-based secured remote server (PureSANs array) with power diagnostics reported weekly from the server provider. The combined power consumption of the server’s production and total flash disk storage was 34 MWh/year. We estimated the data storage energy cost based on the proportion of our laboratory’s data share size (27.39 TB, 2.3% of total server space (1228.8TB)) to be 19 kWh/TB/year which equated to 512 kWh/year. Based on the South Scotland kWh-to-CO_2_e conversion factor (20 g CO_2_e/kWh, Figure 2B), the emissions burden of the laboratory’s data storage amounted to 1.2g CO_2_e per hour or 10.35 kg CO_2_e for the 2023-4 research year. A single year of PhD research generated 1.69TB of data (i.e., raw video files, extracted csv files, python scripts, figures, manuscripts), equivalent to 6.2% of the laboratory’s total data storage space. Thus, the emissions burden of data storage for a single South Scotland PhD student amounted to 0.07g CO_2_e per hour (Figure 1B) or 637.7g CO_2_e for the 2023-4 research year (Figure 1C).

**Figure 2:**
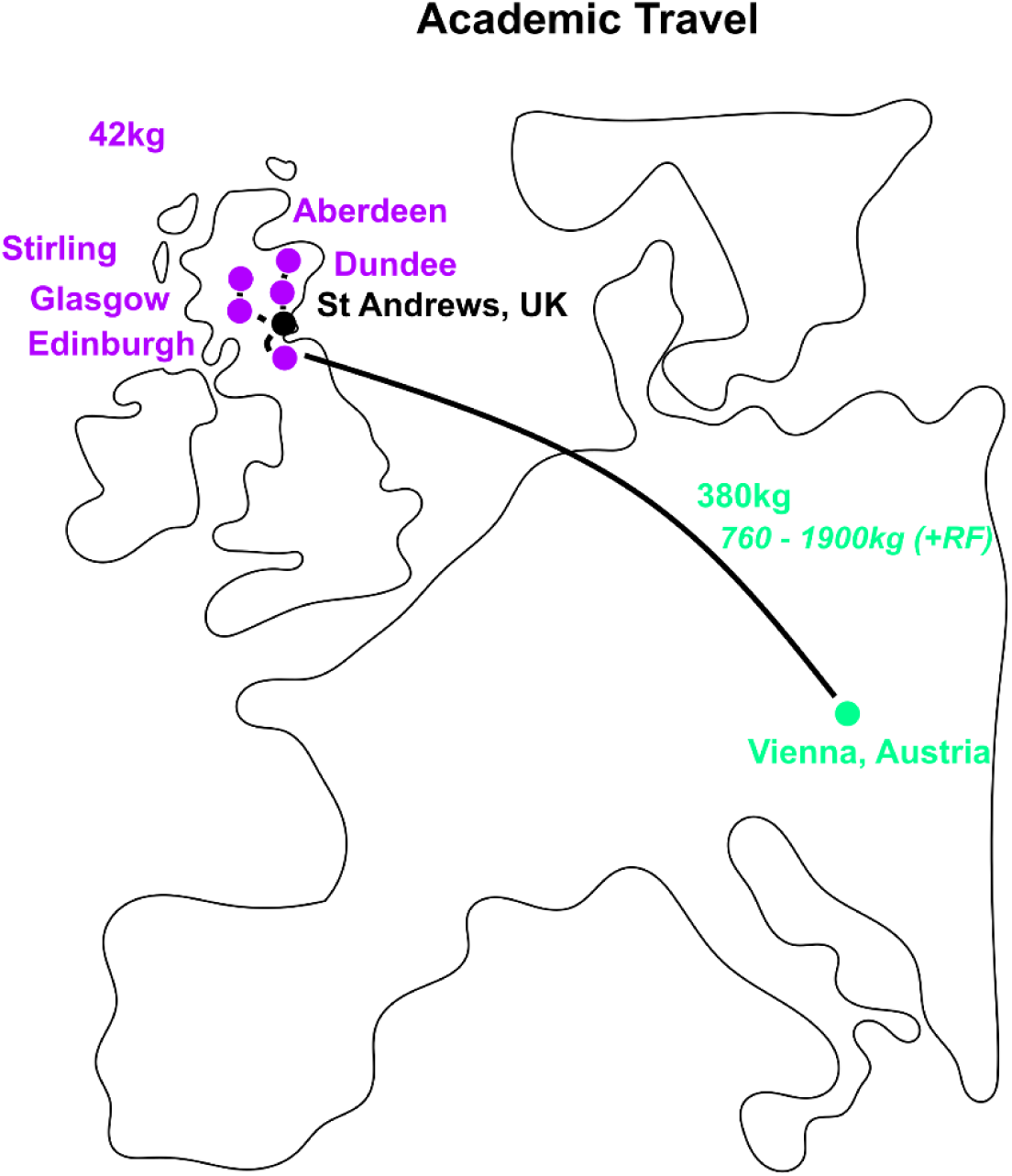
eCO_2_ released via public travel for a PhD researcher 2023-4. Representation of the accrued travel costs of an individual PhD researcher throughout Scotland via national rail (purple) and economy air travel (cyan) without and with (*below*) radiative forcing index factors.

Beyond analysis and storage, data is often communicated, shared, and presented through video streaming applications (i.e., Microsoft Teams) to supervisors, academic staff, and wider academic circles. We found 0.07 kWh was used by our laptop model (Table 1) per 1-hour online meeting where visual and audio material was shared continuously amounting to, on average, 1.42g CO_2_e per instance (Figure 1B). Given our recorded average of four online contact hours per month, we accounted for 68g CO_2_e for the 2023-4 researched year (Figure 1C).

Overall, data storage, analysis, and online presentations produced 1 kg CO_2_e for the 2023-4 research year (1.6x the yearly CO_2_e generated from all recorded experimental work).

#### Disposal

Frontier biological research using genetically modified organisms often requires extensive use of energetically expensive sterilisation and disposal mechanisms, thereby incurring a high indirect energy-related (i.e. scope 2) carbon cost. At the end-stage of the *Drosophila* research life cycle (Figure 1A), high-temperature, high-pressure autoclaves are used to sterilise vials, glassware containing chemical modulatory compounds, and equipment that has contacted genetically modified material. We accounted for the 10-kWh maximum power usage specified by the manufacturer (Table 1), resulting in a maximum average of 202.2g CO_2_e per month for the total laboratory requirement of 700 vials. We found that autoclaving produced a total of 22.7kg CO_2_e for the June 2023 - May 2024 period for the entire laboratory. For a single PhD student (7.4% of total load), autoclave disposal of vials amounted to 15.0g CO_2_e per month (Figure 1B) and a total of 1.7kg CO_2_e (Figure 1C) for the June 2023 - May 2024 period.

#### Persistent equipment

Accompanying the experimental life cycle, equipment for climate control (e.g. air conditioning and incubators) is an essential feature of research facilities. Within our 82.52 m^2^ research space, gas-based heating systems and national grid-supplied electricity for lights were used alongside air conditioning units, incubators, CO_2_ monitoring systems, and fridge/freezer units (Table 1, Figure 1D). The University of St Andrews uses SystemLink™ to dynamically track kWh used for heating and lighting per floor area per building (see Methods). Using SystemLink™ information, we found building maintenance (e.g., heating, lighting) accounted for 36% of total persistent equipment emissions. Surprisingly, incubators accounted for only 5.5% of total persistent equipment CO_2_e emissions, producing on average ∼3.2g eCO_2_ an hour. The persistent CO_2_ monitoring systems yielded an average of 0.2g eCO_2_ an hour. Overall, persistent equipment accounted for the largest proportion of CO_2_e generated per year in our laboratories (212 kg CO_2_e, Figure 1C).

Taken together, one year of *Drosophila* cellular neuroscience research conducted by a PhD student in South Scotland 2023-4 produced 18.15 kg CO_2_e. In particular, across the research life cycle, direct research-related activities generated 3.62 kg scope 1 (i.e., direct release by virgin collection and gas-related heating), 13.84 kg scope 2 (i.e., food generation, experimental work, analytical pipelines, and resource disposal), and 0.71 kg scope 3 (i.e., data storage, distribution, and presentation) CO_2_e emission costs.

### Associated Travel Costs for a Researcher

Aside from conducting research, researchers require travel for training, public engagement, and collaboration. We accounted for the travel-associated carbon emissions for the neuroscience PhD student to provide a case study for non-research related carbon costs of academia (Figure 2). We used ICAO (ICAO, 2024), a free resource for estimating the carbon intensity of travel. ICAO bases carbon emissions measurements exclusively on flight distance, cargo type, plane model, and logged fuel usage per passenger weight. However, it does not include radiative forcing (RF) factors which encapsulate the indirect effects of greenhouse gas emissions caused by contrail formation at higher altitudes during flight (Fuglestvedt *et al*., 2003; Lee *et al*., 2021).

In our case study, travel occurred both internationally via air and locally by rail. Train travel to local Scottish universities for mandatory PhD-related training and research presentation events generated 42 kg CO_2_e (Table 3). Air and associated intermediary travel to the Federation of European Neuroscience (FENS) 2024 conference in Vienna yielded 380 kg CO_2_e (Table 2). Using the comprehensive RF estimation range (1.0 - 4.0) made by Lee et al., (2019), we found that RF may have increased the yearly air travel carbon cost to 760-1900 kg CO_2_e. Thus, the indirect cost of PhD research related to travel amounted to 442 kg CO_2_e (without RF) or 802-1942 kg CO_2_e (with RF) scope 3 emissions. For comparison, the total CO_2_e emissions associated with nine travel instances (without RF) was equivalent to ∼191% of total experiment-related emissions (422 kg versus 2.21 kg) for the 2023-4 research year. In other words, the total travel-related yearly emissions (without RF) were equivalent to the average emissions yield of ∼32 electrophysiology experiments, ∼124 dual colour calcium imaging experiments, ∼334 calcium imaging or ∼618 behavioural optogenetics experiments conducted in South Scotland.

**Table 3:**
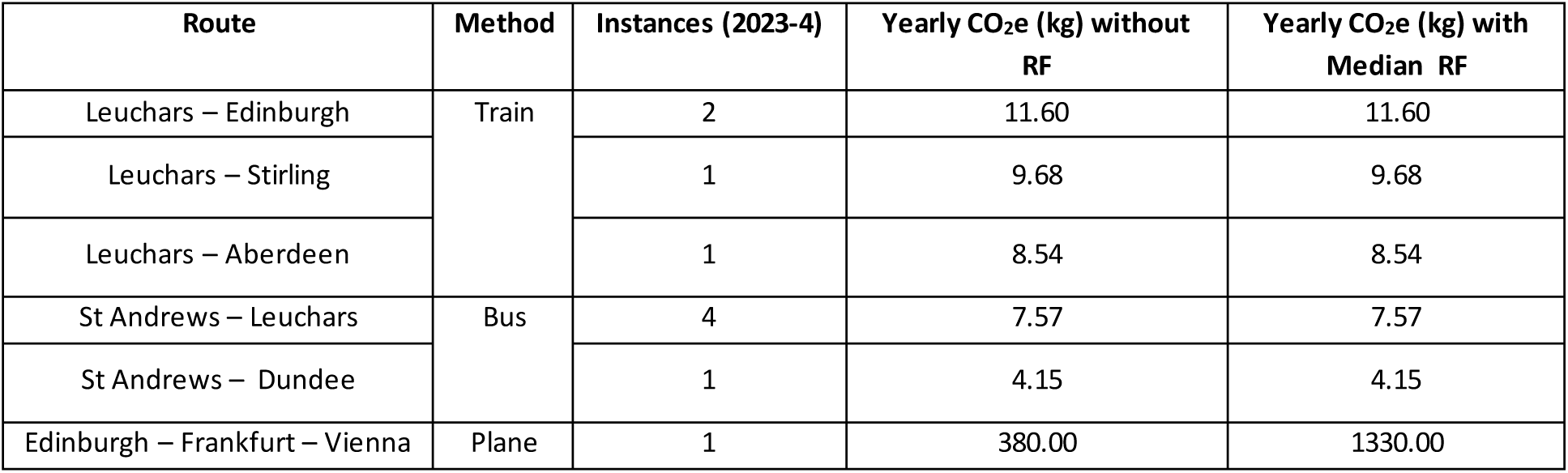
Travel-related CO_2_e Costs for PhD-related Travel for Training and Conferences.

| Route | Method | Instances (2023-4) | Yearly CO<sub>2</sub>e (kg) without RF | Yearly CO<sub>2</sub>e (kg) with Median RF |
| --- | --- | --- | --- | --- |
| Leuchars – Edinburgh | Train | 2 | 11.60 | 11.60 |
| Leuchars – Stirling |  | 1 | 9.68 | 9.68 |
| Leuchars – Aberdeen |  | 1 | 8.54 | 8.54 |
| St Andrews – Leuchars | Bus | 4 | 7.57 | 7.57 |
| St Andrews – Dundee |  | 1 | 4.15 | 4.15 |
| Edinburgh – Frankfurt – Vienna | Plane | 1 | 380.00 | 1330.00 |

### Known Unknowns: The International Cost of *Drosophila* Science

Aside from producing direct emissions (scope 1) or indirect by electricity-based experimental work and building maintenance (scope 2), equipment procurement by laboratories significantly contributes to international emissions (scope 3). Here, we focused on some ubiquitous resources for *Drosophila* research (Figure 3A) and tried to account for their indirect emissions. However, as noted previously (Achten, Almeida and Muys, 2013), companies often reject information requests by deeming transportation routes, times, energy mixtures in manufacturing, and cost as proprietary information. Nevertheless, some information (e.g., manufacture origin, energy mixture in manufacturing, transportation routes) was provided and is sufficient to produce a robust estimate when combined with UK GOV Emission Conversion Data (see Methods).

**Figure 3:**
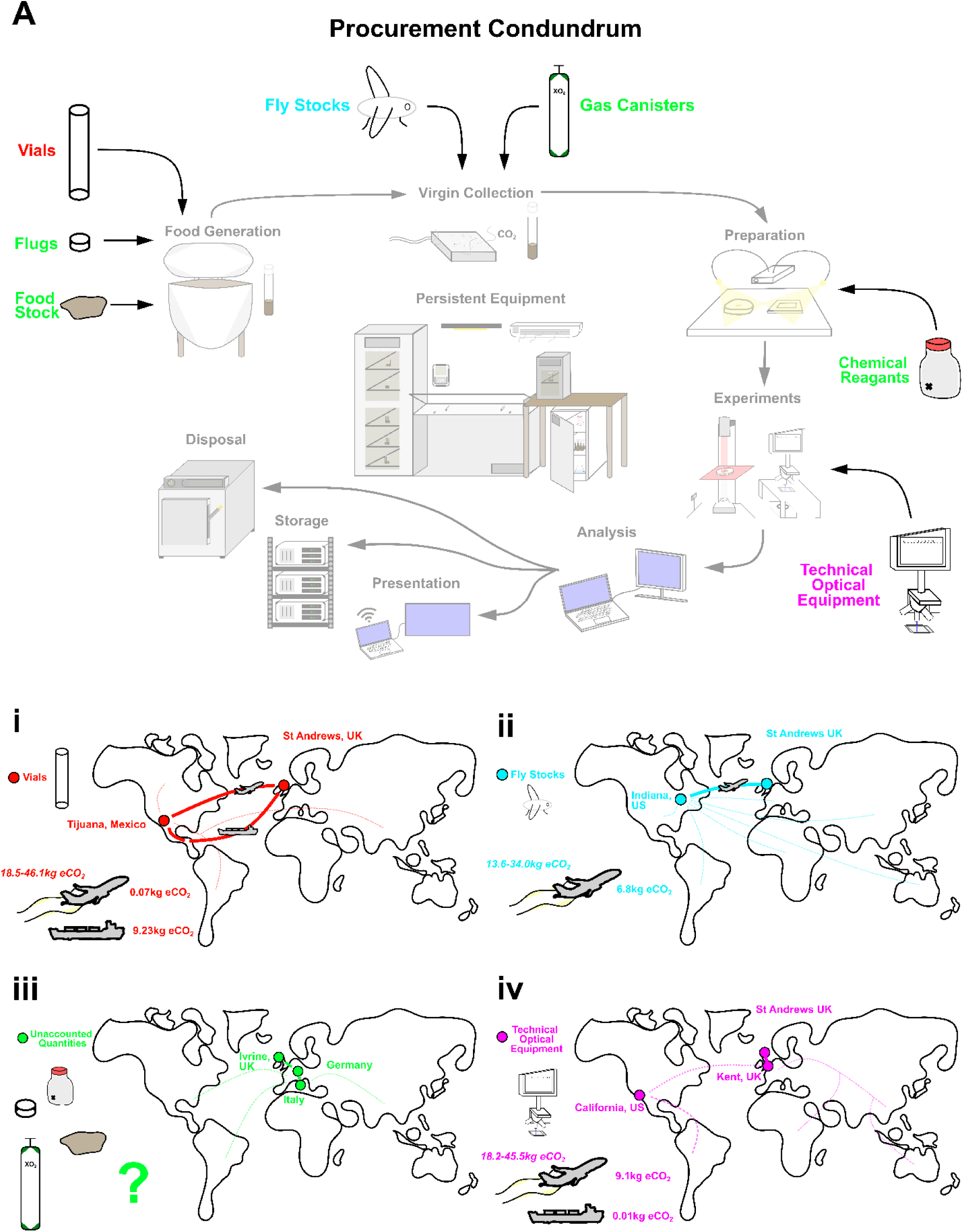
Scope 3 Procurement Emission Estimations for *Drosophila* Biological Research. (**A**) Aside from emissions produced by experimental research and persistent equipment, additional resources also have associated greenhouse emissions at the main stages of research: (**i**) *Drosophila* vial resources are sourced (500 per order) from Mexico through Scientific Laboratory Supplies™, (**ii**) *Drosophila* stocks are mainly sourced from Bloomington Indiana, US with estimate quoted per vial, (**iii**) unaccounted procurement resources: chemical reagents are mainly sourced from Sigma™ with many transport centres across Europe, gas canisters are sourced from BOC™ with Fife-based distributors, flugs and food resources have unknown origin sites (**iv**) optical equipment from CAIRN Research Ltd™ (Kent, UK) or Chroma™ (Irvine, California, US) with one of our orders consisting of four filters and one mirror. Solid lines represent parts of the transport link with eCO_2_ estimates. Dotted lines represent expected but unquantified parts of the transport procurement links upstream in the procurement chain. Radiative factor (1.0 - 4.0) air-travel cost estimates are quoted behind plane images in each panel in italics.

First, we attempted to estimate the cost of sourcing plastic vials for our representative PhD project. Through liaison with Scientific Laboratory Supplies™, we discovered that their basic non-recyclable plastic vials (widely used by *Drosophila* researchers) were sourced from Tijuana, Mexico. However, we were unable to ascertain the exact method of delivery from Mexico to the UK. Thus, we instead used best approximations for distance yielded by using Google’s Distance Matrix, mode of transport assumptions, and UK GOV 2024 conversion factors for CO_2_e for mode of transport (see Methods). If delivery uses high-carbon, faster transport (e.g., air travel with short-distance HGV transfers) we estimated an upper value of 9.23kg CO_2_e per 500 vials for transportation from Tijuana, Mexico to Edinburgh Airport towards the St Andrews, UK. If we incorporate RF estimates (Fuglestvedt *et al*., 2003; Lee *et al*., 2021), air-based procurement of fly vials rises to 18.46 - 46.15kg CO_2_e per 500 vials. However, if delivery uses low-carbon, slower transport (e.g., bulk HGV from Tijuana to Mexico City (Benito Juarez, cargo shipping to London, UK) we estimate a potential 132x reduction in CO_2_e (70g) (Figure 3Ai, Table 4). Given a single PhD student used 624 vials (June 2023 - May 2024), vial acquisition contributed 18.46kg CO_2_e (36.92 - 92.30 kg CO_2_e with RF) if using flight-based procurement or 0.14kg CO_2_e if using cargo-based procurement. For the laboratory as a whole, we acquired 8500 vials in the 2023-4 research period amounting to 156.96kg CO_2_e (313.92 - 784.80 kg CO_2_e with RF) if using flight-based procurement or 1.25kg CO_2_e if using cargo-based procurement.

**Table 4:** Procurement-related CO_2_e costs for Vial and Fly Stocks in 2023-4 Research Year.

| Equipment | Count | Weight per Item (kg) | Total Estimated Travel Distance (HGV + Flight) (km) | CO<sub>2</sub>e (kg) (HGV - Flight) | Total Estimated Travel Distance (HGV + Cargo) (km) | CO<sub>2</sub>e (kg) (HGV – Shipping) |
| --- | --- | --- | --- | --- | --- | --- |
| Vials (500) | 17 | 0.16 | 8464 | 9.23 | 12181 | 0.07 |
| Optic Filters (4 filters, 1 mirror) | 1 | 0.06 | 8287 | 9.11 | 12532 | 0.01 |
| Fly Stocks (4) | 12 | 0.08 | 6317 | 6.79 |  |  |

In addition to vials, *Drosophila* research often relies on importing flies from Bloomington Indiana, US, or alternative stock centers. The time-sensitive nature of transporting live flies necessitates air freight. By using UK GOV 2024 conversion data and Google Matrix’s distance estimations between origin site and final destination, we estimated a production of 6.79 kg CO_2_e (without RF) per vial ordered to St Andrews, UK, generated by cargo flight from Indiana International Airport to Edinburgh Airport followed by HGV travel to St Andrews, UK (Figure 3Aii, Table 4). Incorporating RF, fly procurement may be 13.56 - 33.97kg CO_2_e. Importantly, we could not estimate upstream carbon emissions from research institutions donating *Drosophila* stock to the Indiana-based store nor the precise upkeep costs (though estimates made by this paper could be cross-applied to resources at Bloomingtons at a future date) so the total life-cycle scope 3 cost for *Drosophila* procurement will be larger. Nevertheless, a single PhD student ordered 48 new *Drosophila* stocks from Bloomington’s amounting to 81.48 kg CO_2_e (162.96 - 407.40 kg CO_2_e with RF) for the June 2023 - May 2024 period.

Our research utilised optical equipment that we often continuously source from suppliers like CAIRN Research Ltd™ and producers like Chroma™. While we were unable to acquire information about production and distribution costs from the producers of our cameras, filters, and lenses, CAIRN Research™ did release information about distribution centres and modes of delivery for our 2023-4 items (4 filters, 1 mirror, 0.08kg, Table 4). Our optical orders were sourced from the CAIRN Research Ltd™ facility in Kent, UK by HGV to St Andrews, UK. CAIRN Research Ltd™ procures the optical equipment from the manufacturer intermediate Chroma™ in California, US. Similar to our methodology for vial procurement estimation, we used publicly available cargo route information and UK GOV 2024 conversion factors to estimate low (HGV, cargo ship: 0.01kg CO_2_e per order) and high (HGV, cargo flight from John Wayne to Edinburgh Airport: 9.11kg CO_2_e per order) carbon load procurement routes (Figure 3 Aiv). Incorporating RF, optical procurement may be 15.90 - 43.22kg CO_2_e.

In contrast to our estimations of vial, optical equipment and fly stocks, the lack of publicly available data in chemical reagent supplication (Sigma-Aldrich™, Avantor™ by VMR™, Thermo Fisher Scientific™), gas canister supply (BOC™), or yeast food made any reasonable carbon estimates from those lab dependencies unfruitful. In all, we were unable to discover the prior transportation routes, pre-supplies, or locations of precursor materials earlier in the supply route from any of our chemical suppliers (Figure 3iii). However, we did note down our yearly usage of these materials for future estimation as information becomes more publicly available (Table 5).

**Table 5:** Yearly Procurement Quantity for One PhD Student.

| Equipment | Yearly Used Mass (g) | Quantity | Provider | Serial/Batch Code |
| --- | --- | --- | --- | --- |
| NaCl | 157.50 | < 1 | Sigma Aldrich™ | S9888-1KG |
|  |  | < 1 | VMR™ | 21A144160 |
| KCl | 7.45 | < 1 | Sigma Aldrich™ | P9333-500G |
|  |  | < 1 | VMR™ | 14I170013 |
| CaCl<sub>2</sub> x 2H<sub>2</sub>O | 5.90 | < 1 | VMR™ | 14G090030 |
| MgCl<sub>2</sub> x 6H<sub>2</sub>O | 16.25 | < 1 | Sigma Aldrich™ | M2670-500G |
|  |  | < 1 | VMR™ | 14H18004 |
| TES Buffer | 22.95 | < 1 | Sigma Aldrich™ | T1375-100G |
|  |  | < 1 | Thermo Fisher Scientific™ | B21819.30 |
| Sucrose | 246.50 | < 1 | VMR™ | 201284121 |
| Gas Canister | 39000 | 6 | BOC™ | BOC UN1013 EC 204-696-9 |
| Yeast Food | 6240 |  |  |  |
| Flugs | 624 | < 1 | Scientific Laboratory Supplies™ | FLY1198 |

Overall, total equipment procurement for the 2023-4 year for a single PhD student equated to 81.63 - 542.92kg CO_2_e scope 3 emissions cost. In effect, the median year’s procurement-related emissions (230.65kg CO_2_e) were equivalent to 363x of the total experiment-related emission cost. Altogether, laboratory persistent operations, experimental load, individual academic-related travel, and procurement totalled at < 2502 kg CO_2_e for the 2023-4 year.

### Research Costs Across Space and Time

The carbon intensity of research varies widely depending on the geographical region where it is conducted, largely due to differing energy mixtures across different portions of the UK electricity grid (Figure 4A-B, see Methods). Between June 2023 and May 2024, we estimated that persistent equipment and experimental work together produced 12.6 kg CO_2_e from mains electricity in South Scotland. Applying different regional kWh-to-CO_2_e conversion factors to the total energy consumed by persistent equipment and experimental work yields large differences in estimated CO_2_e emissions. For instance, conducting our research within North Scotland (e.g., Aberdeen) could have reduced total CO_2_e emissions by a factor of 1.87 (to 6.7 kg CO_2_e). In contrast, had our research consumed electricity supplied to South Wales, we would have released 12.3x more CO_2_e (156 kg CO_2_e) than we produced in South Scotland. The largest difference is between North Scotland and South Wales, with on average 23.1x more CO_2_e emissions in South Wales compared to North Scotland.

**Figure 4:**
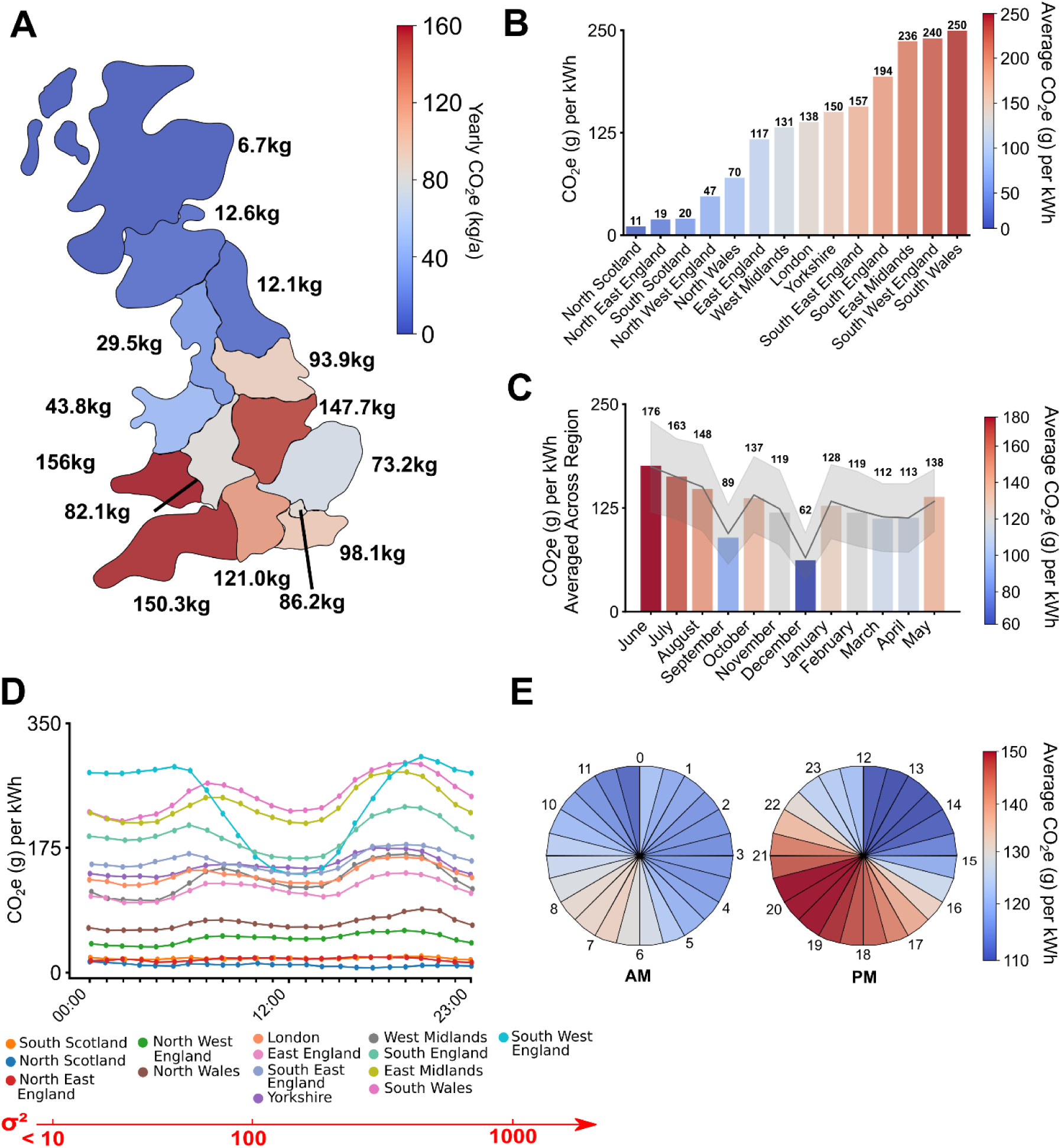
The carbon cost of research by region, month, and time of day in 2023 -4. (**A**) The yearly carbon costs of one researcher and total laboratory costs (total 700 vials per month) in different regions of the UK. (**B**) The average CO_2_e-per-kWh generated from each region of the UK from June 2023 - May 2024. (**C**) The monthly CO_2_e-per-kWh averaged across all regions from June 2023 - May 2024 as recorded by the UK National Grid. The grey line represents the average trend across all regions of the UK while the shaded grey region shows the standard error. (**D**) Variation in the average CO_2_e-per-kWh for different UK regions with time of day, with ranked variance of the regions listed below the time series plot. (**E**) The average CO_2_e-per-kWh for each 30-minute segment on average across regions and months in the UK from June 2023 - May 2024 as recorded by the UK National Grid.

Aside from geography, the month and time that experiments are conducted also affects carbon emissions. For instance, an electrophysiology dataset (N = 10 preparations, 2 hours per prep) collected in South Scotland in December may produce 265 g CO_2_e whereas generating the same dataset in June could result in 2.8x higher carbon emissions (750 g CO_2_e) (Figure 4C).

Similarly, performing experiments at different times of day results in large variations in CO_2_e (Figure 2D, E). For example, performing a dual-colour calcium imaging experiment (1 hour) at 13:00 may produce 11.3 g CO_2_e whereas conducting the same experiment at 19:00 produces on average 36% more CO_2_e (15.5 g). However, temporal variation in CO_2_e-per-kWh is greater in some regions (e.g., Southwest England, σ^2^ = 3505.2 g CO_2_e per kWh) than others (e.g., South Scotland, σ^2^ = 1.9 g CO_2_e per kWh), largely due to differing proportions of renewable resources in regional energy mixtures.

## Discussion

Here, we present the direct and indirect carbon costs of conducting one year of high-intensity, frontier *Drosophila* research using widely used imaging techniques and methods for a single neuroscience PhD student. We show the equivalent carbon emissions (CO_2_e) associated with a variety of biological techniques widely used beyond the field of *Drosophila* research. Alongside scope 1 costs associated with genetic construction, we provide estimates for scope 2 and 3 costs related to research including travel, procurement, and climate control system costs at both the laboratory and individual PhD student level. Additionally, we raise the importance of geography and time when conducting research supplied by national grids that have region-specific resource mixtures (i.e., varying levels of renewable vs non-renewable resources) and varying peaks in energy demand. With variation in mind, we show how the Carbon Intensity API can be used to create region- and time-specific estimations of researchers’ carbon footprints that are more accurate than estimates obtained exclusively using government-reported kwh-to-CO_2_e conversion rates. Such a life cycle assessment exemplifies how doctoral students can make carbon estimations about their research, aiding in the availability of open-access emissions data from the research sector.

Using “cradle-to-grave” life-cycle assessments, the medical and clinical sciences have taken great strides in carbon accounting (e.g., Robinson *et al*., 2018; Maida *et al*., 2024). However, the limited number of individual estimates, inconsistent methodology, and partial attempts at capturing complete life-cycles signals the necessity of increased participation in carbon accounting and consistency in how emissions estimates are calculated (McGain and Naylor, 2014; Purohit, Smith and Hibble, 2021; Valls-Val and Bovea, 2021; Robinson *et al*., 2023). The imminence of key climate targets necessitates efficient individual, collective, and institutional action to mitigate carbon emissions across all sectors. We believe that accurate and publicly available emissions data will be necessary to inform those actions. While a complete dynamic picture of the emissions landscape may be elusive, even a modest increase in emissions data could be readily used to limit and validate large-scale carbon footprint models across disciplines, helping to inform public policy (Ni *et al*., 2018). We suggest that the current paucity of estimations can be partially remedied by engaging researchers at all levels of career progression: a series of individual-led estimation efforts will, together with accurate and consistent carbon emissions estimates, help to capture complete research life cycles even if some researchers can individually account for the carbon produced only by specific life-cycle elements.

The medical and clinical sciences have taken great strides in carbon accounting. Primarily through life-cycle (“cradle-to-grave”) assessments, we now possess both direct and indirect emission estimates for surgical operations (Robinson *et al*., 2023), endoscopy (Maida *et al*., 2024), anaesthesia (McGain *et al*., 2020) medical care infrastructure (McGain and Naylor, 2014), and patient travel (Purohit, Smith and Hibble, 2021). These wide ranging carbon emissions estimates have informed research into effective carbon mitigation strategies at multiple levels of medicine (Montoro *et al*., 2023; Quann *et al*., 2023).

However, the limited number of individual estimates, inconsistent methodology, and partial attempts at capturing complete life-cycles signals the necessity of higher engagement in carbon accounting and consistency in how emissions estimates are calculated (McGain and Naylor, 2014; Purohit, Smith and Hibble, 2021; Valls-Val and Bovea, 2021; Robinson *et al*., 2023). Additionally, multi-sector data could be readily used to limit and validate large-scale carbon footprint models across disciplines (Ni *et al*., 2018). We suggest that the paucity of individual estimations can be partially remedied by engaging researchers at all levels of career progression: a series of individual-led estimation efforts will, together with accurate and consistent carbon emissions estimates, help capture complete research life cycles even if some researchers can only account for the carbon produced by specific life-cycle elements during their research.

Indeed, our emerging understanding of several disciplines’ carbon footprints is often built upon a grassroots approach whereby individual researchers and laboratories contribute their carbon footprint and employ ex-ante and post-hoc mitigation strategies to help the community, for example in astronomy (Martin *et al*., 2022), particle physics (Janot and Blondel, 2022), medical imaging (Picano, 2021), and high-performance computing (Lannelongue, Grealey and Inouye, 2021). Scientists have used these procedural insights to build accreditation programs, such as MyGreenLab, LEAF that seek to inform and reward researchers’ efforts to adopt sustainable research practices (Freese *et al*., 2024).

While work has begun to account for carbon produced by medical neuroimaging (i.e., fMRI) (Souter *et al*., 2023, 2024), to our knowledge, our work is the first to account for cellular neuroscience imaging and *Drosophila*-related research using real-time regional emissions estimates. Thus, our work, methods, and data are intended to plant the seed for further carbon accounting by other disciplines of neuroscience and STEM, across institutions and geographies. Importantly, we envisage PhD-level, individual-driven, lab-focused carbon accounting building on attempts to accurately carbon footprint higher-education institutions (e.g., Valls-Val and Bovea, 2021), to generate an actionable foundation for institutional and government policy for our domain of STEM.

### Methods of Carbon Accounting

Differences in the methods authors use to account for carbon emissions have produced a large variability in estimates (Pandey, Agrawal and Pandey, 2011). Naturally, there are sensible justifications for adopting coarse estimates when more precise estimations (e.g., equipment-specific energy consumption and regional electricity carbon intensity) are unavailable. Complementing Souter *et al* (2023) carbon estimation approach which tracks mains electricity-related emissions using Electricity Maps (https://app.electricitymaps.com/map), we utilised the National Grid-based Carbon Intensity API (https://carbon-intensity.github.io/api-definitions/#carbon-intensity-api-v2-0-0) to demonstrate a more precise method of carbon accounting. However, at the time of writing, region-specific data was restricted to only a few areas (e.g., UK counties and some US states). As climate targets near and international pressure mounts, we predict that demand for APIs like Carbon Intensity will increase, and hope that members of the community will advocate for local governments and private providers to publicise energy-generation mixes so developers can satisfy the need for regional precision in carbon accounting. Focusing on regional precision can enable the research community, including future doctoral students, to disentangle the sources of variability in carbon footprint estimates and thus be better able to formulate effective mitigation strategies.

### Mitigation Strategies

Carbon mitigation is achieved by pre-emptive strategies, which seek to avoid anticipated emissions, and sequestration strategies, which aim to capture the equivalent of previous and unavoidable emissions. Assessment of a carbon footprint is often paired with an equivalent sequestration comparative (Elli *et al*., 2024). However, sufficient carbon sequestration is a complex demand with factors like tree species, the plantation ecosystem, climate change, biodiversity, and soil ecology all contributing to varying mitigation success (Woodward *et al*., 2009; Abbas *et al*., 2017; Vacek, Vacek and Cukor, 2023). In addition, the rise of business models around reforestation commonly incentivise short-term sequestration followed by biomass extraction which renders the mitigation strategy effectively redundant. Alternative sequestration methods (e.g., carbon-capture-storage) are often too expensive for individuals and university institutions to participate in the short term to reach 2030-50 climate targets, especially considering the financial trade-off made against more solvent renewable - led investment for emission mitigation (Sgouridis *et al*., 2019). While post-release mitigation is necessary to combat decades of over release of greenhouse gases, the complementary, more robust and immediately accessible method of mitigation is pre-emptively reducing activity emissions and procurement demands, leaving sequestration efforts to capture historic and unavoidable emissions via large private and government-led initiatives. Like most human decisions, assessing and knowing the quantity associated with a harm or benefit can be an additive, sufficient motivating factor for action (Zhang *et al*., 2019). For example, our work revealed unexpectedly high energy consumption by persistent climate control systems due to aging, inefficient equipment. This has motivated us to acquire more energy efficient equipment with effects we can quantify and report consistently in the future. Alongside persistent equipment, the amount of direct emissions generated by *Drosophila* sorting for genetic crosses far exceeded our expectations. Considering the wide usage of *Drosophila* due to their genetic versatility, the scale of CO_2_ release by the *Drosophila* community will be considerable. According to FlyBase’s Fly Lab list (https://wiki.flybase.org/wiki/FlyBase:Fly_Lab_List), there were 2,074 active Drosophila laboratories as of December 2024. Considering we estimated 4kg CO_2_e per research, if we conservatively project an average *Drosophila* laboratory size of 3 researchers per lab, this may amount to 25,000kg CO_2_e per year which, according to ICAO’s flight emission calculator, is equivalent to 40 return economy trips from Edinburgh to New York’s JFK airport. Thus, we recommend that researchers investigate and invest in less carbon-intensive methods where appropriate, for *Drosophila* researchers this could take the form of anaesthetisation by cooling when powered by renewable resources. More broadly, by following the method and standards reported here, doctoral students in other scientific disciplines can expose other systemic, high-emission scientific practices, informing and motivating transitions to more sustainable methods.

Efforts to account for greenhouse gas emissions across disciplines have historically used non-regional, general kWh-to-CO_2_e conversion rates that have contributed to wide variation in reported emissions. Here, we clearly demonstrate the impact of geography and time when calculating emissions estimates. The fact that changing our lab’s location within the UK could increase our annual research carbon footprint by an order of magnitude raises questions about the best geographical location for high-energy research during the transition to 100% renewable energy. Importantly, large-scale relocation of high-energy research to a currently greener region would evidently create diminishing returns if energy supply and/or storage were inelastic. Notwithstanding, medium-term large-scale decarbonisation efforts of national grids could be complemented by short-term considerations of regional variation in energy mixtures when flexibility in the location of research or facilities exists.

While our calculations assumed electricity from the National Grid, many institutions have invested considerable effort into in-house generation of renewable, low emission energy (Adedeji, Akinlabi and Madushele, 2019; Vaziri, Rezaee and Monirian, 2020). This does, however, raise ethical questions concerning the link between financial privilege and the ability to invest in energy solutions: care must be taken to encourage creative, collaborative emissions mitigation strategies that are not restricted to select institutions. Indeed, even the production of low-emissions energy is not immediately net-neutral; emissions are expended in construction resources, maintaining infrastructure, and transport emissions aimed to provide “greener” in-house supply (e.g., from solar power: Wu *et al*., 2021). Such ethical ramifications are analogous to treatment-versus-waste questions reported in medical carbon accounting (Thiel and Richie, 2022). While we cannot resolve the ethical question, we do offer effective suggestions on research-level pre-emptive mitigation routes applicable to all institutions regardless of geographic, economic, or social constraints.

#### The Procurement Conundrum

We used a bottom-up approach to estimate the scope 3 footprint associated with material procurement (“cradle-to-gate”) for our research. A significant effort has been invested by the medical and adjacent community to account for material procurement costs at both an individual and institutional level (Clabeaux *et al*., 2020; Montoro *et al*., 2023; Quann *et al*., 2023; Maida *et al*., 2024). Like other carbon accounts (McGain *et al*., 2020; Maida *et al*., 2024; Souter *et al*., 2024), we used government-reported statistics on fuel conversion alongside reasonable approximations of the mode and route of transport. In the case of vial and fly provision, we acquired information about product assembly, location of distribution centres, weight of shipped products and amount of recyclable materials both via public means and by interacting with procurer sustainability representatives. However, necessary data concerning product manufacture, assembly, and distribution were often held back due to proprietary concerns. Specifically, we were unable to ascertain: i) the exact mode of transport used in sub-parts of the distribution chain, ii) the complete distribution route, iii) the location, number, or energy-demands of prior routes in the distribution chain (i.e., raw resource procurement), iv) the energy mix of the manufacturer factories, or v) manufacturer-level mitigation strategies. While we can make robust travel cost estimations if we know the start point of the distribution chain, the precision of the account is significantly impaired by a lack of industry information and transparency (Stenzel and Waichman, 2023). In some instances (e.g., gas canisters, chemical reagents, cotton flugs to seal *Drosophila* inside vials), we were unable to acquire any information related to procurement due to a lack of response to information requests. Nevertheless, the majority of distributors we contacted (3 of 5) were approachable, enthusiastic, receptive, and open to information requests, demonstrating how overcoming the initial energy barrier of reaching out to procurers can reveal novel and utile information. For instance, alongside learning about the procurement routes from the Bloomington *Drosophila* stock centre, we also learned the total volume of *Drosophila* distribution to laboratories worldwide in 2023. The future must see both procurement institutions and researchers attempt to track, estimate, and record carbon-related procurement information so we can better focus and evaluate emission mitigation targets. Researchers are capable of only so much when provided with incomplete information. Institutions must report on upstream procurement (e.g., routes, methods, and loads from primary producers) as well as downstream procurement information (e.g., routes, methods, and loads to consumers). The manufacturers and distributors that are first-movers in publicly accessible and accurate emission-related data will not just gain a significant foot in the carbon net-neutral landscape but will enable researchers to act as changemakers by making active choices to support more sustainable distributors and manufacturers, creating a race-to-openness by suppliers and distributors in and beyond academia.

Procurers and intermediate agents in the scientific resource supply chain often have highly useful information that can be easily accessed by direct contact. The highly connected, dynamic and demanding landscape for procurement often results in agents involved in the supply chain having access to precise, accurate, and highly useful information. However, the sheer volume of data or the lack of obvious public uses for supply chain meta-data (e.g., number of orders, user’s location, frequency of orders) often mean that it is not made publicly accessible, increasing the energy barrier for widespread, individual-led carbon accounting. For instance, direct contact with Bloomington *Drosophila* Stock Centre yielded disclosure of 2023 outflow information for domestic and worldwide *Drosophila* laboratories (e.g., 171,426 vials, 17 vials per order, single orders ranging from 1-200 vials, 50% of US to 50% international orders). This exemplifies how individual-led action can yield meta-data from procurers that can help to refine and scale up estimates of carbon footprints from the researcher to the community and international level.

Moving forward, we hope that community-driven action in neuroscience and adjacent disciplines will foster a culture of openness amongst our industry partners, kick-starting a race-to-openness on emissions data. The initiative shown by Scientific Laboratory Supplies and other companies demonstrates how researchers and suppliers can work together to inform more sustainable research practices. Indeed, the European Union (EU) have already begun legislating for carbon accounting at a higher industry level which will continue to see EU producers and distributors release energy and CO_2_e-related information in the coming years (Dinh, Husmann and Melloni, 2023; Hummel and Jobst, 2024). Supporting these efforts is a rise in initiatives derisking and aiding stakeholders (e.g., multi-stakeholder initiatives) to effectively meet legislative targets (Romito, Vurro and Pogutz, 2024). At the individual level, researchers are beginning to calculate the carbon footprint of their research, generating robust datasets that enable effective policy implementation and accountability at the institutional level. Supporting the next generation of carbon literate researchers will multiply the impact of academic institutions as change-makers in the race towards net-neutral, informing procurement decisions that provide a market incentive for data openness among equipment producers and distributors.

#### The Future of Research CO_2_e

To reach our climate targets, large-scale grid decarbonisation efforts must be complemented by technological and social innovation to improve the efficiency and accessibility of renewable resources, and increasing carbon literacy, with its attendant changes in individual behaviour (Sithole *et al*., 2016). These actions may be hampered by institutional and individual complacency, redirection of decarbonisation funding to inefficient streams, and the rising dominance of high-energy industries (e.g., complexification of AI, blockchain mining for cryptocurrency) (Truby *et al*., 2022; Cowls *et al*., 2023). Even without such barriers, careful planning is necessary to avoid increases to scope 3 emissions when expanding renewable energy infrastructure (e.g., mineral mining for batteries, artificial intelligent-required computer chips, renewable machinery, system-wide heat-pump adoption) with consideration for equipment longevity and recycling (Jackson, Meinke and Chandramohan, 2024).

Within academia, climate targets will undoubtedly be challenged by increasingly energy-intensive research methods and social resistance to short-term transition costs. Fortunately, high emission sectors including medicine (McGain and Naylor, 2014) and agriculture (Jebari *et al*., 2024) are leading the way in carbon accounting for informed emissions mitigation, paving the way for other disciplines to interrogate their emissions landscapes, to develop strategies to meet institutional, national, and international emissions reduction targets (Zhao *et al*., 2024). We believe that this effort will be most effective if individuals and institutions work together to collect accurate emissions data for efficient, equitable, and community-driven transitions to sustainable research practice.

Our work shows how individual researchers in any discipline can accurately and efficiently account for greenhouse equivalent emissions. Mirroring medical and clinical researchers, we have commenced an open-source carbon data store for *Drosophila* neuroscience research. Alongside our meta-analysis reported here, we point researchers to efforts at emissions tracking at all levels of research (Loyarte-López, Barral and Morla, 2020; Mariette *et al*., 2022) that are aiding our understanding of how different research techniques, geographies, travel demands, and scientific practices affect carbon emissions. We encourage researchers at all levels to participate in building a codex of carbon accounts that will empower the scientific community to lead the way as a positive social force for climate accountability.

## Methods

### Power Estimation

The power usage of experimental and laboratory equipment was determined using RS Pro Energy Metres (Stock No: 178-5370; Model No. PM01; Stock No. 123-3230). The power demands of equipment associated with experiments (e.g., calcium imaging) was recorded for 1-3 instances of mock experiments by using the power metres interlocked from wall mains supply. The power of persistent equipment (e.g., incubators, fridges) was recorded over a 24-72 hr period in February-March 2023. We estimated the power usage of our analysis using the hardware analysis and monitoring software HiWinFO during analysis of experimental-related data (e.g., traces, Python-based visualisation) (https://www.techspot.com/downloads/5245-hwinfo64.html). Clamp metres (Stock No. 123-3230) were used to measure energy from air conditioning units. The high-powered equipment for food generation and autoclaves precluded direct measurement for safety reasons, thus we resorted to maximum power calculations based on the equipment’s industry reported metrics. The power usage of our laboratory spaces was monitored through direct building power metres reported by SystemsLink™ and subsequently applied to our research based on floor area (electricity: 71.48kWh/m^2^; gas: 170.14kWh/m^2^).

### Direct CO_2_ Release from Virgin Collection

The collection of *Drosophila* virgins for genetic crosses necessitates anaesthesia by direct application of CO_2_ to sort flies by sex and fertility status. *Drosophila* adults are initially anaesthetised within their vial using a gas gun, then tipped onto a CO_2_ pad where CO_2_ is intermittently released to arrest movement. We estimated the direct release of CO_2_ during virgin collection from a gas gun (SLS FlyStuff Blowgun FLY1042) through a graduated 100ml glass gas syringe (9L per minute) and from a CO_2_ pad (SLS FlyStuff Flypad Standard FLY1208) using a fluid displacement method. We timed the application of CO_2_ during virgin collecting amongst three researchers to arrive at the average CO_2_ released per vial during virgin collection. We scaled up the measurement for heavy genetic screening laboratories based on reported vial load for two and three stage genetic constructs.

### Power-to-eCO_2_ Conversion

We utilised the open-source, freely available National Grid ESO’s carbon intensity regional Carbon Intensity API to convert the power usage of equipment associated with experiments and persistent equipment to a valid equivalent carbon dioxide (CO_2_e) estimation. For the 2023-4 period, count per month of experiments were recorded and translated into CO_2_e estimations using the API. Specifically, we took the mean of the first two weeks of each month (June 2023-4) to calculate our monthly-associated research eCO_2_ emissions. The detailed methodology of the regional carbon intensity API is available on GitHub (https://carbon-intensity.github.io/api-definitions/#carbon-intensity-api-v2-0-0). Code for data extraction from the API, processing, and visualisation is available in our repository.

### Travel Cost & Procurement Estimation

Estimates of CO_2_e travel costs were made using a combination of the UK 2024 government’s reported conversion factors (https://www.gov.uk/government/publications/greenhouse-gas-reporting-conversion-factors-2024), self-reporting by ScotRail (https://www.scotrail.co.uk/carbon-calculator) and the ICAO Carbon Emissions Calculator (ICAO) (https://www.icao.int/environmental-protection/Carbonoffset/Pages/default.aspx). The methodology for each report is detailed through provided links. We used Google’s Distance Matrix API to calculate the distance of journeys.

We estimated the CO_2_e emissions associated with equipment transport during procurement based on the reported site of manufacture. Non-recyclable vials (FLY1312, Scientific Laboratory Supplies) are manufactured and assembled in Tijuana, Mexico. We aimed to estimate the range of associated CO_2_e emissions for procurement from Tijuana to St Andrews based on distance matrix API calculations and reported destinations for travel. We assumed vial exports delivered by air freight came directly from Tijuana International Airport to Edinburgh Airport. We assumed cargo-based exports came from the main Mexican export port, Manzanillo Port. We assumed direct travel between the reported final destination of manufacture and the final procurement destination. *Drosophila* fly stocks were procured from Bloomington Indiana. We assumed air freight from Indianapolis International Airport to Edinburgh International Airport. We assumed that vehicle transport of both vials and flies used an average HGV per the UK government’s reported emissions values.

## Acknowledgments

We would like to acknowledge and thank Lucy Moore and Megan Gray from Scientific Laboratory Supplies (SLS), Christopher Warren (CAIRN Research Ltd™) for their efforts to push suppliers on information requests on behalf of the authors alongside. Further, we would like to thank our laboratory technician Amy Dorward for her help with understanding our gas canister supply and advice in persistent energy measurements, and St Andrews’ Information Officer Dean Drew for his assistance in understanding data centre power costs. We thank M. Pankratz at the University of Bonn for constructive discussions and comments on drafts of this manuscript. Further, we thank Dr Jason Jacques for their consultation about the NESO national grid. A.B. sincerely thanks Marcus Bischoff for access to laboratory facilities, and Marcus Bischoff and Jochen Kursawe for supervision and comments on the manuscript.

This project was made possible by funding from the Scotland’s Future Series (University of St Andrews) to WVS, an Industrial CASE PhD studentship (UKRI Biotechnology and Biological Sciences Research Council (BBSRC) grant number BB/T00875X/1) to WVS, a World Leading PhD studentship (University of St Andrews) to AB and M. Bischoff, a Global PhD studentship (University of St Andrews and University of Cologne) to RS, SRP and M. Gather. This work was additionally supported in part by two Collaborative Research Grants awarded jointly by the Global Office of the University of St Andrews and The Halle Institute for Global Research at Emory University (SRP and A. Prinz), and The Global Office of the University of St Andrews and the University of Bonn (SRP and M. Pankratz), and by a Neurophotonics pump-priming Grant from the RS Macdonald Foundation (SRP and M. Gather).

## Data Availability

The data is available upon reasonable request to the corresponding author.

## Author’s Contributions

S.R.P. and W.V.S. conceived the study. W.V.S. and A.B. determined scope 1 emissions. W.V.S. and R.S. performed energy usage estimates. W.V.S. and S.R.P. engaged with suppliers and distributors to estimate scope 3 emissions. W.V.S. constructed analytical pipelines and drafted the figures and manuscript. A.B. provided assistance with emissions calculations and revised the manuscript. W.V.S. and A.B. finalised the figures. S.R.P. supervised the project.

## References

1. Abbas, F. et al. (2017) ‘Agroforestry: a sustainable environmental practice for carbon sequestration under the climate change scenarios—a review’, Environmental Science and Pollution Research, 24(12), pp. 11177–11191. Available at: 10.1007/s11356-017-8687-0.

2. Achten, W.M.J., Almeida, J. and Muys, B. (2013) ‘Carbon footprint of science: More than flying’, Ecological Indicators, 34, pp. 352–355. Available at: 10.1016/j.ecolind.2013.05.025.

3. Adedeji, P.A., Akinlabi, S. and Madushele, N. (2019) ‘Powering the future university campuses: a mini-review of feasible sources’, Procedia Manufacturing, 35, pp. 3–8. Available at: 10.1016/j.promfg.2019.05.003.

4. Budennyy, S.A. et al. (2022) ‘eco2AI: Carbon Emissions Tracking of Machine Learning Models as the First Step Towards Sustainable AI’, Doklady Mathematics, 106(S1), pp. S118–S128. Available at: 10.1134/S1064562422060230.

5. Clabeaux, R. et al. (2020) ‘Assessing the carbon footprint of a university campus using a life cycle assessment approach’, Journal of Cleaner Production, 273, p. 122600. Available at: 10.1016/j.jclepro.2020.122600.

6. Cowls, J. et al. (2023) ‘The AI gambit: leveraging artificial intelligence to combat climate change— opportunities, challenges, and recommendations’, AI & SOCIETY, 38(1), pp. 283–307. Available at: 10.1007/s00146-021-01294-x.

7. Dinh, T., Husmann, A. and Melloni, G. (2023) ‘Corporate Sustainability Reporting in Europe: A Scoping Review’, Accounting in Europe, 20(1), pp. 1–29. Available at: 10.1080/17449480.2022.2149345.

8. Elli, L. et al. (2024) ‘The carbon cost of inappropriate endoscopy’, Gastrointestinal Endoscopy, 99(2), pp. 137–145.e3. Available at: 10.1016/j.gie.2023.08.018.

9. Farley, M. and Nicolet, B.P. (2023) ‘Re-use of laboratory utensils reduces CO2 equivalent footprint and running costs’, PLOS ONE. Edited by A. Jahanger, 18(4), p. e0283697. Available at: 10.1371/journal.pone.0283697.

10. Fenner, A.E. et al. (2018) ‘The carbon footprint of buildings: A review of methodologies and applications’, Renewable and Sustainable Energy Reviews, 94, pp. 1142–1152. Available at: 10.1016/j.rser.2018.07.012.

11. Freese, T. et al. (2024) ‘The relevance of sustainable laboratory practices’, RSC Sustainability, 2(5), pp. 1300–1336. Available at: 10.1039/D4SU00056K.

12. Fuglestvedt, J.S. et al. (2003) ‘Metrics of Climate Change: Assessing Radiative Forcing and Emission Indices’, Climatic Change, 58(3), pp. 267–331. Available at: 10.1023/A:1023905326842.

13. Gordon, I.O. et al. (2021) ‘Life Cycle Greenhouse Gas Emissions of Gastrointestinal Biopsies in a Surgical Pathology Laboratory’, American Journal of Clinical Pathology, 156(4), pp. 540–549. Available at: 10.1093/ajcp/aqab021.

14. Grealey, J., et al. (2022) ‘The Carbon Footprint of Bioinformatics’, Molecular Biology and Evolution. Edited by S. Kumar, 39(3), p. msac034. Available at: 10.1093/molbev/msac034.

15. Häusser, M. (2014) ‘Optogenetics: the age of light’, Nature Methods, 11(10), pp. 1012–1014. Available at: 10.1038/nmeth.3111.

16. Heitler, W.J. (2007) ‘DataView: A Tutorial Tool for Data Analysis. Template-based Spike Sorting and Frequency Analysis’, *Journal of undergraduate neuroscience education: JUNE: a publication of FUN*, Faculty for Undergraduate Neuroscience, 6(1), pp. A1–7.

17. Higher Education Statistics Agency (HESA) (2024). Available from: https://www.hesa.ac.uk/news/08-08-2024/sb269-higher-education-student-statistics

18. Hummel, K. and Jobst, D. (2024) ‘An Overview of Corporate Sustainability Reporting Legislation in the European Union’, Accounting in Europe, 21(3), pp. 320–355. Available at: 10.1080/17449480.2024.2312145.

19. ICAO (2024). Available from: https://applications.icao.int/icec/Methodology%20ICAO%20Carbon%20Emissions%20Calculator_v13_Final.pdf

20. Jackson, L., Meinke, C. and Chandramohan, R. (2024) ‘Challenges in the Battery Raw Materials Supply Chain: Achieving Decarbonisation from a Mineral Extraction Perspective’, *Mining*, Metallurgy & Exploration, 41(5), pp. 2683–2692. Available at: 10.1007/s42461-024-01070-7.

21. Janot, P. and Blondel, A. (2022) ‘The carbon footprint of proposed $${\mathrm{e}}^+{\mathrm{e}}^-$$ Higgs factories’, The European Physical Journal Plus, 137(10), p. 1122. Available at: 10.1140/epjp/s13360-022-03319-w.

22. Jebari, A. et al. (2024) ‘Feasibility of mitigation measures for agricultural greenhouse gas emissions in the UK. A systematic review’, Agronomy for Sustainable Development, 44(1), p. 2. Available at: 10.1007/s13593-023-00938-0.

23. Knot, H.J. et al. (2005) ‘Twenty years of calcium imaging: cell physiology to dye for’, Molecular Interventions, 5(2), pp. 112–127. Available at: 10.1124/mi.5.2.8.

24. Lakovaara (1969) ‘Malt as a culture medium for Drosophila species’, Drosophila Inf. Serv, 44, p. p.128.

25. Lannelongue, L., Grealey, J. and Inouye, M. (2021) ‘Green Algorithms: Quantifying the Carbon Footprint of Computation’, Advanced Science, 8(12), p. 2100707. Available at: 10.1002/advs.202100707.

26. Lee, D.S. et al. (2021) ‘The contribution of global aviation to anthropogenic climate forcing for 2000 to 2018’, Atmospheric Environment, 244, p. 117834. Available at: 10.1016/j.atmosenv.2020.117834.

27. Loyarte-López, E., Barral, M. and Morla, J.C. (2020) ‘Methodology for Carbon Footprint Calculation Towards Sustainable Innovation in Intangible Assets’, Sustainability, 12(4), p. 1629. Available at: 10.3390/su12041629.

28. Maida, M. et al. (2024) ‘Green endoscopy, one step toward a sustainable future: Literature review’, Endoscopy International Open, 12(08), pp. E968–E980. Available at: 10.1055/a-2303-8621.

29. Mariette, J. et al. (2022) ‘An open-source tool to assess the carbon footprint of research’, Environmental Research: Infrastructure and Sustainability, 2(3), p. 035008. Available at: 10.1088/2634-4505/ac84a4.

30. Marley, R. and Baines, R.A. (2011) ‘Dissection of Third-Instar *Drosophila* Larvae for Electrophysiological Recording from Neurons’, Cold Spring Harbor Protocols, 2011(9), p. pdb.prot065656. Available at: 10.1101/pdb.prot065656.

31. Martin, P. et al. (2022) ‘A comprehensive assessment of the carbon footprint of an astronomical institute’, Nature Astronomy, 6(11), pp. 1219–1222. Available at: 10.1038/s41550-022-01771-3.

32. Mathis, A. et al. (2018) ‘DeepLabCut: markerless pose estimation of user-defined body parts with deep learning’, Nature Neuroscience, 21(9), pp. 1281–1289. Available at: 10.1038/s41593-018-0209-y.

33. Mazzetto, A.M., Falconer, S. and Ledgard, S. (2022) ‘Mapping the carbon footprint of milk production from cattle: A systematic review’, Journal of Dairy Science, 105(12), pp. 9713–9725. Available at: 10.3168/jds.2022-22117.

34. McGain, F. et al. (2020) ‘Environmental sustainability in anaesthesia and critical care’, British Journal of Anaesthesia, 125(5), pp. 680–692. Available at: 10.1016/j.bja.2020.06.055.

35. McGain, F. and Naylor, C. (2014) ‘Environmental sustainability in hospitals – a systematic review and research agenda’, Journal of Health Services Research & Policy, 19(4), pp. 245–252. Available at: 10.1177/1355819614534836.

36. Montoro, J. et al. (2023) ‘Impact of Asthma Inhalers on Global Climate: A Systematic Review of Their Carbon Footprint and Clinical Outcomes in Spain’, Journal of Investigational Allergy and Clinical Immunology, 33(4), pp. 250–262. Available at: 10.18176/jiaci.0887.

37. Ni, K. et al. (2018) ‘Carbon Footprint Modeling of a Clinical Lab’, Energies, 11(11), p. 3105. Available at: 10.3390/en11113105.

38. OECD (2019) *Education at a Glance 2019: OECD Indicators*. OECD (Education at a Glance). Available at: 10.1787/f8d7880d-en.

39. Ozlu, E. et al. (2022) ‘Carbon Footprint Management by Agricultural Practices’, Biology, 11(10), p. 1453. Available at: 10.3390/biology11101453.

40. Panagiotopoulou, V.C., Stavropoulos, P. and Chryssolouris, G. (2022) ‘A critical review on the environmental impact of manufacturing: a holistic perspective’, The International Journal of Advanced Manufacturing Technology, 118(1–2), pp. 603–625. Available at: 10.1007/s00170-021-07980-w.

41. Pandey, D., Agrawal, M. and Pandey, J.S. (2011) ‘Carbon footprint: current methods of estimation’, Environmental Monitoring and Assessment, 178(1–4), pp. 135–160. Available at: 10.1007/s10661-010-1678-y.

42. Picano, E. (2021) ‘Environmental sustainability of medical imaging’, Acta Cardiologica, 76(10), pp. 1124–1128. Available at: 10.1080/00015385.2020.1815985.

43. Purohit, A., Smith, J. and Hibble, A. (2021) ‘Does telemedicine reduce the carbon footprint of healthcare? A systematic review’, Future Healthcare Journal, 8(1), pp. e85–e91. Available at: 10.7861/fhj.2020-0080.

44. Quann, N. et al. (2023) ‘Reducing the carbon footprint of research: experience from the NightLife study’, BMJ Open, 13(4), p. e070200. Available at: 10.1136/bmjopen-2022-070200.

45. Rae, C.L. et al. (2022) ‘Climate crisis and ecological emergency: Why they concern (neuro)scientists, and what we can do’, Brain and Neuroscience Advances, 6, p. 23982128221075430. Available at: 10.1177/23982128221075430.

46. Raugei, M., Kamran, M. and Hutchinson, A. (2020) ‘A Prospective Net Energy and Environmental Life-Cycle Assessment of the UK Electricity Grid’, Energies, 13(9), p. 2207. Available at: 10.3390/en13092207.

47. Robinson, O.J. et al. (2018) ‘Towards a universal carbon footprint standard: A case study of carbon management at universities’, Journal of Cleaner Production, 172, pp. 4435–4455. Available at: 10.1016/j.jclepro.2017.02.147.

48. Robinson, P. et al. (2023) ‘The carbon footprint of surgical operations: a systematic review update’, The Annals of The Royal College of Surgeons of England, 105(8), pp. 692–708. Available at: 10.1308/rcsann.2023.0057.

49. Romito, S., Vurro, C. and Pogutz, S. (2024) ‘Joining multi-stakeholder initiatives to fight climate change: The environmental impact of corporate participation in the Science Based Targets initiative’, Business Strategy and the Environment, 33(4), pp. 2817–2831. Available at: 10.1002/bse.3639.

50. Scanziani, M. and Häusser, M. (2009) ‘Electrophysiology in the age of light’, Nature, 461(7266), pp. 930–939. Available at: 10.1038/nature08540.

51. Schell, B.R. and Bruns, N. (2024) ‘Lab sustainability programs LEAF and My Green Lab®: impact, user experience & suitability’, RSC Sustainability, 2(11), pp. 3383–3396. Available at: 10.1039/D4SU00387J.

52. Sgouridis, S. et al. (2019) ‘Comparative net energy analysis of renewable electricity and carbon capture and storage’, Nature Energy, 4(6), pp. 456–465. Available at: 10.1038/s41560-019-0365-7.

53. Sithole, H. et al. (2016) ‘Developing an optimal electricity generation mix for the UK 2050 future’, Energy, 100, pp. 363–373. Available at: 10.1016/j.energy.2016.01.077.

54. Souter, N.E. et al. (2023) ‘Ten recommendations for reducing the carbon footprint of research computing in human neuroimaging’, Imaging Neuroscience, 1, pp. 1–15. Available at: 10.1162/imag_a_00043.

55. Souter, N.E. et al. (2024) ‘Measuring and reducing the carbon footprint of FMRI preprocessing in FMRIPREP’, Human Brain Mapping, 45(12), p. e70003. Available at: 10.1002/hbm.70003.

56. Stenzel, A. and Waichman, I. (2023) ‘Supply-chain data sharing for scope 3 emissions’, npj Climate Action, 2(1), p. 7. Available at: 10.1038/s44168-023-00032-x.

57. Tamburrini, G. (2022) ‘The AI Carbon Footprint and Responsibilities of AI Scientists’, Philosophies, 7(1), p. 4. Available at: 10.3390/philosophies7010004.

58. Thiel, C. and Richie, C. (2022) ‘Carbon Emissions from Overuse of U.S. Health Care: Medical and Ethical Problems’, Hastings Center Report, 52(4), pp. 10–16. Available at: 10.1002/hast.1404.

59. Thiel, C.L. et al. (2017) ‘Cataract surgery and environmental sustainability: Waste and lifecycle assessment of phacoemulsification at a private healthcare facility’, Journal of Cataract and Refractive Surgery, 43(11), pp. 1391–1398. Available at: 10.1016/j.jcrs.2017.08.017.

60. Tinevez, J.-Y. et al. (2017) ‘TrackMate: An open and extensible platform for single-particle tracking’, Methods, 115, pp. 80–90. Available at: 10.1016/j.ymeth.2016.09.016.

61. Truby, J. et al. (2022) ‘Blockchain, climate damage, and death: Policy interventions to reduce the carbon emissions, mortality, and net-zero implications of non-fungible tokens and Bitcoin’, Energy Research & Social Science, 88, p. 102499. Available at: 10.1016/j.erss.2022.102499.

62. Vacek, Z., Vacek, S. and Cukor, J. (2023) ‘European forests under global climate change: Review of tree growth processes, crises and management strategies’, Journal of Environmental Management, 332, p. 117353. Available at: 10.1016/j.jenvman.2023.117353.

63. Valls-Val, K. and Bovea, M.D. (2021) ‘Carbon footprint in Higher Education Institutions: a literature review and prospects for future research’, Clean Technologies and Environmental Policy, 23(9), pp. 2523–2542. Available at: 10.1007/s10098-021-02180-2.

64. Vaziri, S.M., Rezaee, B. and Monirian, M.A. (2020) ‘Utilizing renewable energy sources efficiently in hospitals using demand dispatch’, Renewable Energy, 151, pp. 551–562. Available at: 10.1016/j.renene.2019.11.053.

65. Woodward, F.I. et al. (2009) ‘Biological Approaches to Global Environment Change Mitigation and Remediation’, Current Biology, 19(14), pp. R615–R623. Available at: 10.1016/j.cub.2009.06.012.

66. Wu, X. et al. (2021) ‘Is solar power renewable and carbon-neutral: Evidence from a pilot solar tower plant in China under a systems view’, Renewable and Sustainable Energy Reviews, 138, p. 110655. Available at: 10.1016/j.rser.2020.110655.

67. Zhang, Y. et al. (2019) ‘Quantified or nonquantified: How quantification affects consumers’ motivation in goal pursuit’, Journal of Consumer Behaviour, 18(2), pp. 120–134. Available at: 10.1002/cb.1752.

68. Zhao, A. et al. (2024) ‘High-ambition climate action in all sectors can achieve a 65% greenhouse gas emissions reduction in the United States by 2035’, npj Climate Action, 3(1), p. 63. Available at: 10.1038/s44168-024-00145-x.

